# Overcoming bottlenecks for in vitro synthesis and initial structural insight of ice nucleating protein InaZ

**DOI:** 10.1101/334987

**Authors:** Irina V. Novikova, Swarup China, James E. Evans

## Abstract

Unlike inorganic or other synthetic alternatives, ice nucleating proteins (INPs) remain the most efficient ice nuclei today. Their potential applications in cryo-preservation, biomedicine, food industry and in the modulation of climate are widespread. Nevertheless, over several decades, cell-based recombinant methods have experienced multiple difficulties expressing these large proteins in full-length and in necessary yields while retaining functionality. As a result, our understanding of the structure and ice nucleation mechanism for this class of proteins is incomplete, and, most importantly, the full extent of possible applications unrealized. Using a wheat-germ cell-free expression pipeline, we successfully expressed and purified full-length ice nucleating protein InaZ from *Pseudomonas syringae*, known as a model INP. High protein yield and solubility has been achieved using this system. Ice nucleation experiments inside a dynamic environmental scanning electron microscope (ESEM) confirmed that the produced InaZ products remain functional. Preliminary structural assessments of these proteins using Transmission Electron Microscopy (TEM) showed experimental evidence for their structural organization as fibrils. We believe that the current platform will be suitable for expressing other INPs of interest and can be further employed as new engineering system either for industrial or scientific needs.

## 1. Introduction

Ice nucleating proteins, INPs, belong to a rare class of proteins that evolved a highly unique ability to enhance ice nucleation in the supercooled water at sub-zero temperatures. These proteins reside in the outer cell membrane of a few select ice-nucleation-active (Ina+) bacterial strains, which include *Pseudomonas* and *Erwinia* species [1,2]. These bacterial species have been widely detected in air and cloud water, suggesting key roles in the atmospheric glaciation and cloud evolution [3-6] while the presence of these organisms on leaf surfaces causes significant frost injuries to plants [7,8].

The most active Ina+ strain known to date is *Pseudomonas syringae* [1]. The first ice-nucleating protein described at the sequence level originated from that strain and contains exactly 1,200 amino acids (aa) [9]. This protein was designated as InaZ and remains the most-characterized INP today. It is divided into three main domains: N-terminal (1-175 aa), central (176-1151 aa) and C-terminal (1152-1200 aa). While the N-terminal domain is responsible for anchoring to the outer membrane of the cell and transmembrane transport [10], the central domain was found to be at the core of the ice nucleation mechanism. This central region is the largest with an unusual highly repetitive architecture where individual 8-residue blocks (112 of them) organize in 16-residue repeats, which are a part of higher-order 48-residue arrangements [9]. Deletion experiments of the various portions of this domain did not pinpoint to an individual subregion responsible for the ice activity and rather suggested a quantitative contribution of individual repeats to the ice formation mechanism [9,11].

Experimental *in situ* radiation studies have shown that the size of active ice-nucleating functional protein assemblies is variable and depends on the nucleation temperature [12]. At lower temperature range (−12 °C to −13 °C), the minimum mass of active complexes was found to correspond to a single protein of 150 kDa. Upon the increase in the ice nucleation temperature, the mass of the complex increases log-linearly and achieves the maximum of 8000 kDa at −2 °C, which corresponds roughly to a multimeric assembly of 53 InaZ proteins. Many additional experimental and modelling efforts have also shown that the multimerization of InaZ or its fragments seem to be critical for its nucleation performance [13-15]. However, questions still remain regarding the full-length structure of InaZ and the detailed mechanism of its action.

To date, the majority of all preliminary characterization has been done on natively-purified InaZ from the cell membranes of *Pseudomonas syringae*, where quantities are generally not sufficient or pure enough for structural work. Several cell-based recombinant expression trials on closely related INPs yielded protein within inclusion bodies when over-expressed and required extra solubilization [16,17]. Additionally, cell-based recombinant expression of InaZ in *E. coli* has only been successful in production of soluble fragments up to 576 residues, yet the fragments did not display any detectable nucleation activity [18]. Finally, commercial peptide biosynthesis services are typically limited to 200 amino acids or less.

A new approach, which can synthesize and purify InaZ protein while simultaneously retaining its functionality, is needed in order to answer many of the open questions in the field of InaZ mechanism, structure, and kinetics. In this work, we demonstrate that, by using wheat germ cell-free protein expression technology, we can successfully express and purify InaZ recombinantly while maintaining the ice nucleation function. We also provide preliminary structural characterization of this protein by Transmission Electron Microscopy.

## 2. Results and Discussion

### 2.1. InaZ amino acid sequence corrected

Due to previous unsuccessful trials to express and purify full-length InaZ in *E. coli* [18], we decided to utilize a wheat germ cell-free expression system, a plasmid-based cell-free platform for high yield synthesis of proteins which displays high solubility rates and quick turnaround timeframes [19-21].

During our cloning efforts transferring the commercially acquired InaZ gene from the host vector to our desired plasmid for the cell-free protein synthesis, we discovered that all generated clones have a number of mutations located in the central region of the gene that change several amino acids compared to the published sequence (Fig. S1, Supplementary file: DNA and amino acid sequence information for InaZ). To eliminate the possibility that these mutations were the result of our cloning efforts, we sequenced the Addgene acquired host plasmid and the same mutations were detected. Through discussions with the original research group who donated the plasmid to the Addgene repository, we were informed that no mutations were introduced before submission and that the plasmid contains the wild type sequence of InaZ from *Pseudomonas syringae*. Surprisingly, the last sequence record for InaZ dated back to 1993 (GenBank: X03035.1) and, despite the extensive interest in this protein, the sequence has not been updated since. Interestingly, the newly-discovered nucleotide mutations in the Addgene plasmid result in amino acid changes that are in better consensus with the rest of the central region repetitive segments (Fig. S1). This suggests that the amino acid discrepancy is the most likely the result of sequencing errors in early days.

### 2.2. Using the wheat germ cell-free format, full-length InaZ is expressed in soluble form and in high yield

Initially, we generated two template plasmids where one contained a full-length InaZ gene and the second had a truncated version with the N-terminal domain removed. The latter InaZ fragment lacking the first 175 amino acids, is referred herein as InaZΔN. Based on early reports, we posited that the N-terminal domain does not play role in ice nucleation and that its removal could potentially eliminate a predicted hydrophobic domain that might hinder solubility. Both plasmids had a 3xFLAG purification tag positioned on the N-terminal of the gene. During our cell-free expression trials, InaZ and InaZΔN proteins were both well expressed and were found soluble. Unfortunately, we were unable to purify those using a 3xFLAG tag (data not shown) - a procedure that was well optimized in house for many other proteins [19]. We thought that a highly hydrophilic 3XFLAG tag (pI 3.6) might potentially interact with the central domain of InaZ and thus prevent the tag’s proper binding to the separation matrix. Thus, we next introduced a more neutral V5 epitope tag (pI 5.71) in both plasmid constructs. Once again, both V5-fused constructs were successfully expressed but the purification procedures failed for both (data not shown). It therefore seemed that tags located at the N-terminus were inaccessible. Our next strategy was to re-position the 3xFLAG tag on the C-terminus of the gene and to conduct additional purification trials (Fig.1). The cell-free synthesis produced soluble InaZ and InaZΔN at approximately equal yield. It is noteworthy that both full-length InaZ (123kDa with the tag) and InaZΔN (103 kDa with the tag) displayed retarded mobility on the SDS-PAGE, displaying higher molecular mass of >150 kDa. The same unique behavior of INPs using PAGE has been reported previously [17,22], suggesting that the cell-free synthesized proteins have retained such unique properties.

**Fig. 1.**
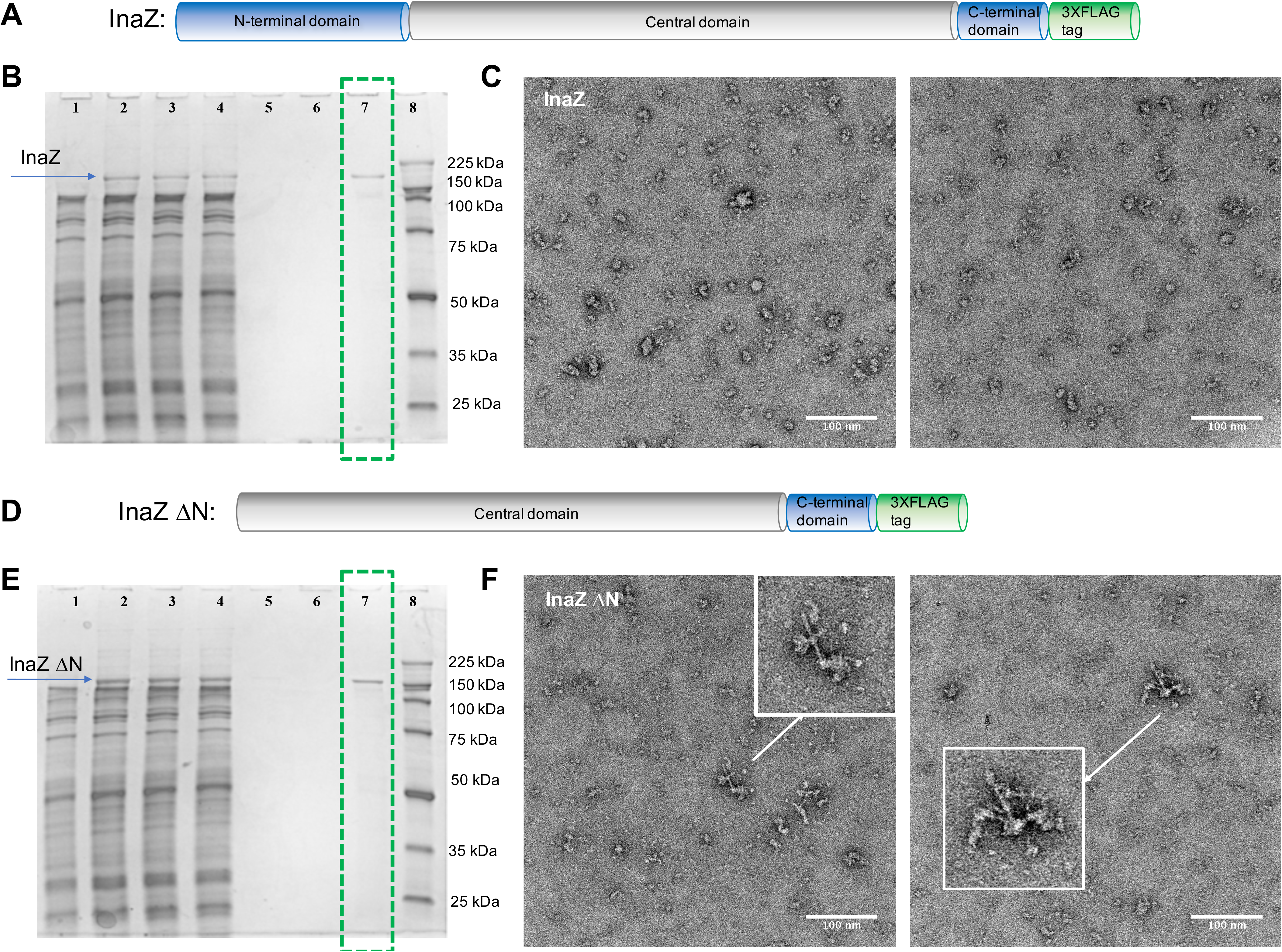
Cell-free synthesis of InaZ proteins and preliminary negative stain TEM characterization. A. Schematic of InaZ. B. SDS-PAGE gel showing expression and purification of full-length InaZ. Lane 1, control expression with no plasmid (negative control); Lane 2, crude mixture; Lane 3, soluble fraction; Lane 4, flow-through; Lane 5, elution fraction 1; Lane 6, elution fraction 2; Lane 7, final concentrated product; Lane 8, MW markers. C. TEM images of purified InaZ product (highlighted in green dashed box). D. Schematic of InaZΔN. E. SDS-PAGE gel showing expression and purification of InaZΔN. Lane annotations are the same as in (B). F. TEM micrographs of purified InaZΔN product. The zoom-in TEM images of rod-like structures are denoted with white arrows.

Despite low recovery of InaZ proteins using 3XFLAG, the final purified proteins were of good purity and amounts, suitable for a preliminary characterization using Transmission Electron Microscopy in combination with negative staining (Fig.1). As clearly seen from a few select TEM images, both InaZ and InaZΔN proteins lacked any defined oligomeric state. Quite heterogeneous population of complexes, which range from small protein assemblies to higher molecular weight protein aggregates, are clearly observed. Interestingly, a frequent occurrence of rod-like protein organization was manifested in the case of InaZΔN (Fig. 1F). No such architectures were found in the full-length InaZ (Fig. 1C). One possible explanation is that the presence of a relatively hydrophobic N-terminal domain interfered with the formation of such rod-like assemblies.

According to previous computational modeling studies the 16 amino acid repeats (in the central domain) are believed to form loops, which are further organized into a β-helix reminiscent of a tube or filament for which a InaZ monomer would theoretically have dimensions of 2 nm in width and around 30 nm in length [14, 23, 24]. The rod-like fibril structures observed here with TEM for InaZΔN are very similar to the theoretical model both in shape and dimensions and represent the first direct experimental evidence of their existence. However, most of the InaZΔN proteins did not exist in this fibril form, which may indicate that the absence of cell membrane lipids or membrane curvature offers more protein conformational sampling in a three-dimensional space. Overall, the obtained results on the cell-free expressed InaZ were consistent with previous scientific reports done on native InaZ which suggested a heterogeneous multimeric nature for this protein [12,13,15]. The high-order of InaZ assembly/organization, where tags are hidden, can explain why tag-based purification approaches largely failed. For reference, in crude mixture, we note that 6 ml translation reaction generated 130 μg of protein in the case of InaZ and 144 μg of protein in the case of InaZΔN. The purification procedure was only slightly more efficient where 2.5% of product was recovered for both constructs.

### 2.3. Complexation of InaZ with liposomes to improve the purification procedure

While the purification is not needed for industrial needs, structural/mechanistic characterization requires access to the purified protein. The wheat germ cell-free synthesis is an “open-format” system where translation mixture can also be supplemented with liposomes to create a potentially stabilizing environment for newly synthesized hydrophobic regions of protein and at the same time providing alternate means for protein purification via density centrifugation [25]. Due to the presence of a hydrophobic N-terminal domain in the full-length InaZ, we attempted cell-free protein expression in the presence of liposomes (Fig. S2). This format yielded 375 μg of InaZ (from 2.5 ml bilayer-dialysis reaction) where 12% of the protein product (complexed with liposome) were purified from the rest of the crude protein mixture by centrifugation. This protein product is referred to as InaZ-liposome herein. The liposome complexation improved the total yield of purified protein and even though the full protein recovery was not achieved, the procedure looked promising. We attributed the low yield to the lack of defined multipass transmembrane domains in the N-terminal domain of the protein. As such we decided to engineer a fusion protein TM-InaZ where the full-length InaZ is coupled with a 20 amino acid artificial transmembrane region [26] on its N-terminus to improve insertion into liposomes. The liposome-assisted cell-free expression and purification of TM-InaZ was repeated, and the purification efficiency had dramatically improved to 75% (Fig. 2B, lane 5). The experiment (both synthesis and purification protocols) was conducted along with the negative control preparation where no DNA template for InaZ synthesis was added to the translation mixture (Fig. 2B, lanes 1 and 2). In comparison with this control, an additional band at around 100 kDa molecular weight in the final purified InaZ sample originates from nonspecific protein absorption to the liposome mixtures.

**Fig. 2.**
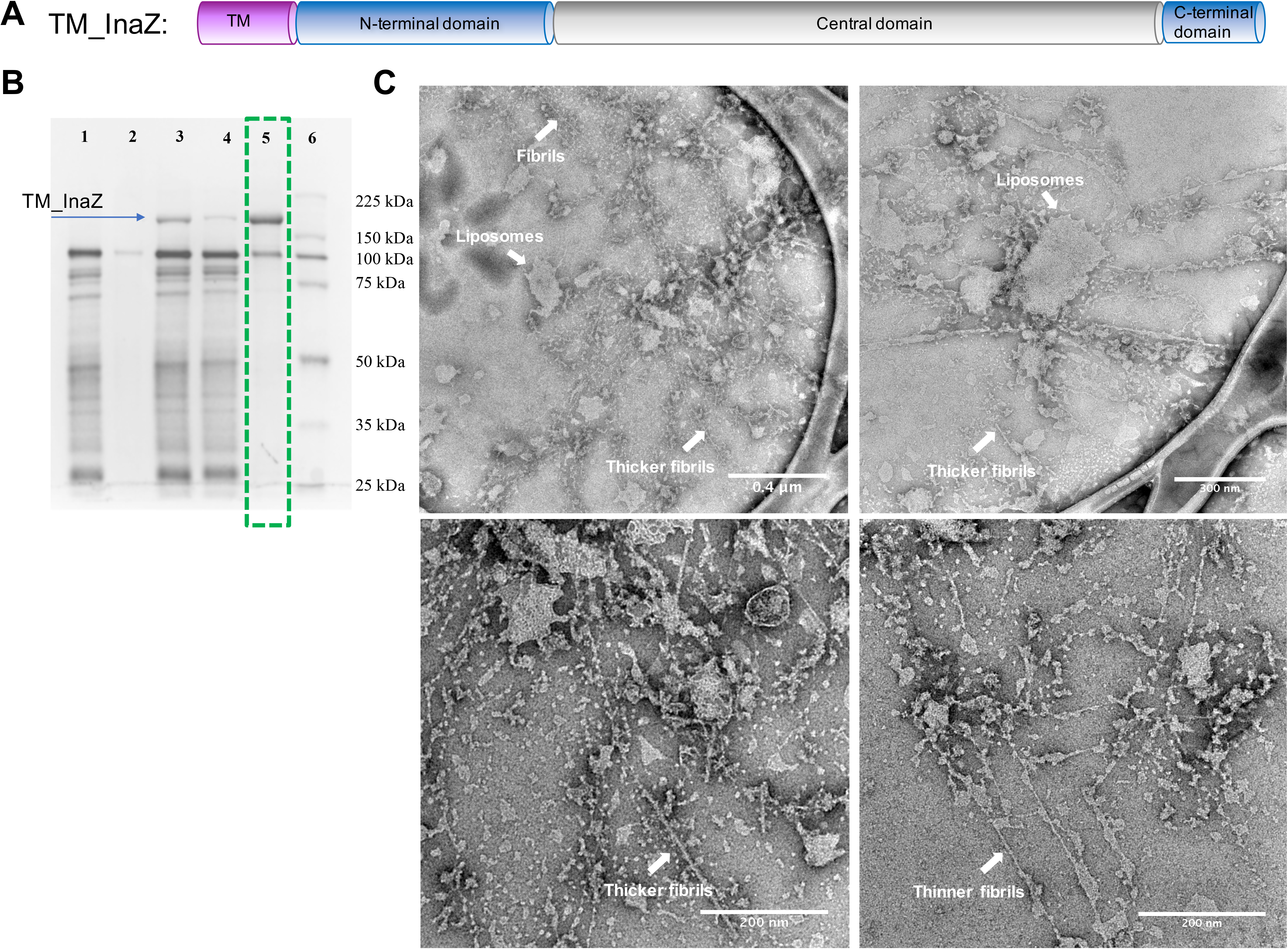
Cell-free synthesis and TEM characterization of TM-InaZ in the presence of liposomes. A. Schematic of TM-InaZ protein construct. B. SDS-PAGE of TM-InaZ synthesis and liposome-assisted purification scheme. Lane 1, control expression with no plasmid; Lane 2, re-suspended pellet (final product of control samples); Lane 3, crude mixture; Lane 4, supernatant (or flow-through); Lane 5, re-suspended pellet (final product); Lane 6, MW markers. C. Negative stain TEM micrographs of TM-InaZ product, complexed with liposomes.

### 2.4. Preliminary insights in InaZ structure

Despite of initial characterization of InaZ occurring in 1985 [9], it is very surprising that experimental structure and the structures of other discovered INP today still remain unsolved and the detailed mechanism for their ice nucleation remains incomplete ever since. Native properties of InaZ such as large size and tendency to aggregation represent a challenge in gaining the access to full-length and functional INP. The same factors hindered prior attempts for recombinant cell-based expression and purification efforts. For example, while a prior paper described the biophysical characterization of 240-residue InaZ fragment produced via cell-based expression and showed that InaZ exists (most likely) as an elongated monomer and is composed mainly of β-sheets and random coil, the same fragment did not display any detectable nucleation activity [18].

During the purification procedure, we noticed that liposomes containing InaZ proteins have changed appearance and texture. Their volume doubled making them look fluffier, and at the same time they became slightly sticky unlike control sample with empty liposome. While the liposome association provides obstacles for the detailed protein characterization by TEM, preliminary structural characterization of TM-InaZ-liposome samples was attempted by negative stain. It became apparent that, in TM-InaZ-liposome sample, there is a prevalence of fibrous net-like interwoven assemblies with liposomes (Fig. 2C) that absent in the control sample (Fig. S3). Two types of fibrous assemblies are apparent: (i) thinner fibrils with the approximate diameter of 2-3 nm, and (ii) thicker fibrils with the approximate diameter of 4-6 nm. Thicker fibrils emerge as straight rods of variable length with no apparent curvature. Even though the thinner fibrils appear mainly straight, a slight curvature is evident suggesting a slightly flexible association. Prior computational modeling on the close homolog of InaZ from *P. borealis* predicted that INPs can dimerize by bridging the active sites. Such theoretical dimeric architecture, where one β-helix tube-like INP monomer binds another monomer unit side-by-side in parallel would result in a flat structure with the width doubled from 2 nm to 4 nm [14]. The dimerization interface involves the most conserved residues of the protein and hypothetically significantly increases the stability and the “ice-nucleation active” surface area of the structure. While the length of thicker fibrils is variable and exceeds the length of the proposed dimer on its own, the width dimension differences between thinner and thicker fibrils are consistent with the proposed dimerization mechanism.

### 2.5. Cell-free produced InaZ is functional

To confirm that these proteins were functional, we conducted ice nucleation experiments using dynamic ESEM to determine the onset of early ice nucleation events, or if not functional the failure to nucleate prior to water saturation. The ESEM images clearly demonstrated that whenever InaZ protein was present in the sample, ice nucleation occurs prior to water saturation (Fig. 3). At 250 K, the onset of nucleation occurred at 113% relative humidity with respect to ice (RH_ice_) for InaZΔN, at 109 % RH_ice_ for InaZ-liposome and at 115% RH_ice_ for TM-InaZ-liposome. The obtained RHice values for cell-free produced proteins were in excellent agreement with the values previously recorded for SNOMAX (a commercial brand name for lysed *P. syringae* cell fractions), which were in 110 to 120% range [27]. Based on these results, and consistent with previous reports, it seems that the N-terminal 175 amino acids do not impact ice nucleation significantly.

**Fig. 3.**
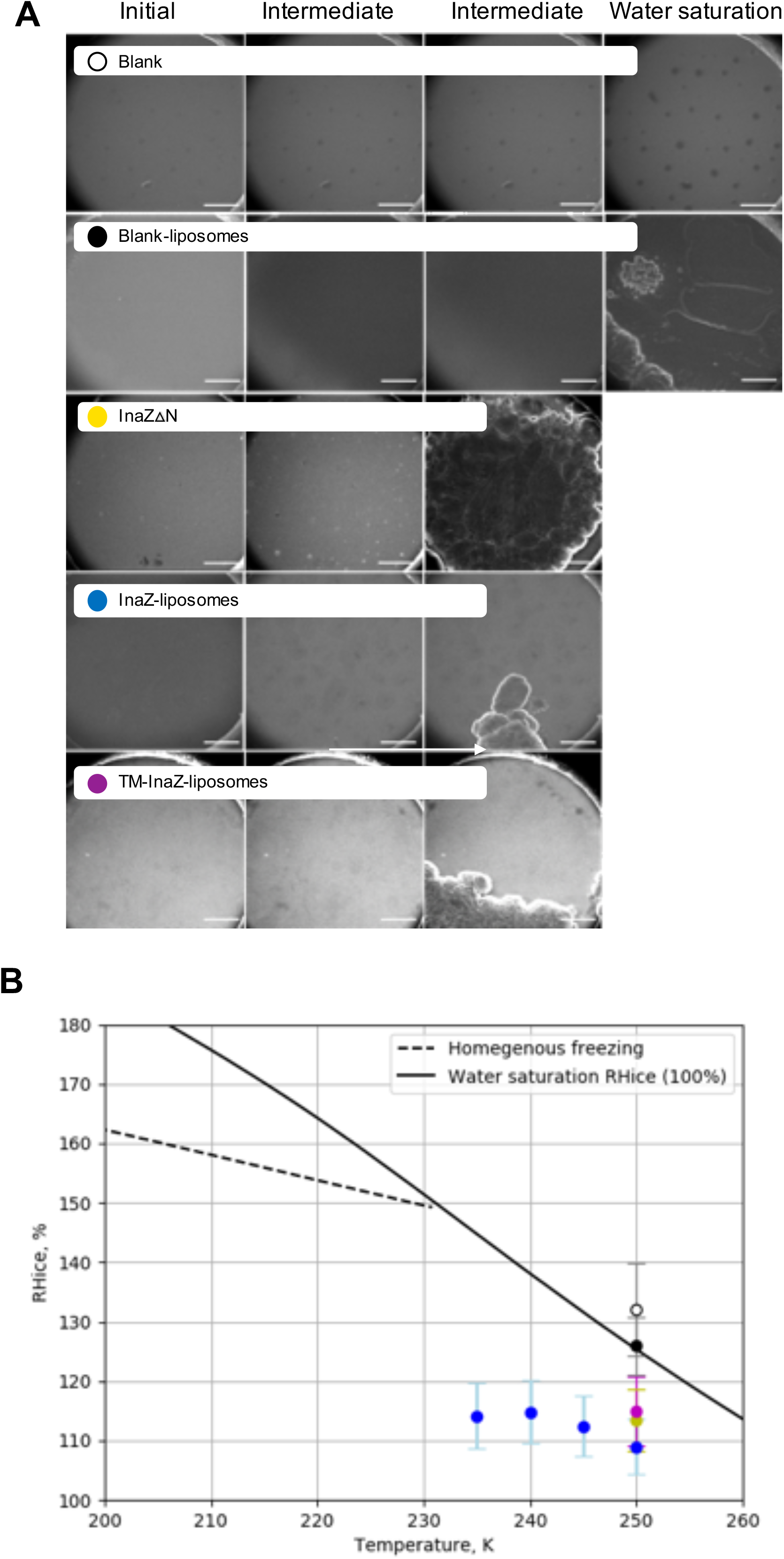
Monitoring ice formation on the sample droplets using dynamic ESEM. A. Recorded ESEM images show initial, intermediate and water saturation conditions. The white scale bar corresponds to 100 microns. B. The mean relative humidity with respect to ice (RH_ice_) for the nucleation event has been calculated. Droplets from InaZ-liposome samples were investigated at multiple temperatures to derive RH_ice_. The error bars were derived from experimental uncertainties.

## 3. Implications and future directions

Bacterial ice nucleation potential has been studied extensively due to their roles in precipitation and frost injury to plants. For industry, the bacterium *Pseudomonas syringae* that possesses InaZ on its outer layer has been utilized and is known as the best ice-inducing agent for decades in the snow-making industry [28]. Partially purified InaZ from native sources has also been more recently demonstrated to perform reliable freeze-thaw valving in the microfluidic devices, providing also a reduction in cost and design complexity of these elements [29]. Unachievable by other cell-based recombinant methods, our results demonstrate the successful synthesis of full-length InaZ protein in a soluble form using the wheat-germ cell-free pipeline. As there was no loss in InaZ functionality, we expect that other INP proteins can be successfully expressed using the same approach. The flexibility of cell-free expression also portends the ability to construct and express synthetic InaZ analogs with desired properties in the future. For example, adjusting the number of repeats may allow tuning of freezing conditions. Alternatively, rapid cell-free expression in combination with mutational analysis can provide future help in understanding the ice nucleation mechanism as a function of the protein molecular properties.

We also provided preliminary structural insight in the native state of InaZ. Observed fibril architectures are consistent with its previously theorized monomer fold and dimerization potential. While InaZ structural heterogeneity remains a bottleneck to overcome, future experiments can explore various constructs to see if one or several might stabilize a single oligomeric state and improve chances for high-resolution structural analysis. Overall, the successful expression of InaZ in a soluble form using the cell-free pipeline described here opens many new avenues for potential research on this environmentally and industrially relevant class of proteins.

## 4. Material and methods

### 4.1. The InaZ vector construction for cell-free protein synthesis

A host plasmid containing a major portion of InaZ gene (aas 1- 1179) was obtained from Addgene plasmid repository, #26759 [30]. Using PCR, the gene was amplified and extended to include additional DNA sequence to encode purification tags via primers. All PCR reactions were carried out using the Q5 Hot Start High-Fidelity 2X Master Mix from New England Biolabs. All DNA primers were purchased from Invitrogen. The template vector was pEU (pEU-MCS-TEV-HIS-C1) from CellFree Sciences. Obtained PCR products, such as linearized pEU template and the gene insert, were first gel-purified using Zymoclean Gel DNA recovery kit from Zymo Research and subjected to Gibson Assembly. The Gibson Assembly Master Mix from New England Biolabs was used as recommended. The assembled product has been used to transform an *E. coli*, which was later plated on a carbenicillin-containing agar plate and grown at 37°C overnight. Several individual colonies were selected and grown overnight in LB medium, supplemented with carbenicillin. Plasmids were extracted using QIAGEN Plasmid Mini Kit and further sent for sequencing to MCLAB. Successful clones were later grown in larger quantities of LB broth (with carbenicillin) and harvested using the QIAGEN Plasmid Maxi kit.

### 4.2. Cell-free protein expression and 3XFLAG purification of InaZ proteins

Protein synthesis was carried out at MAXI-scale (6 ml translation volume) in a bilayer format using Wheat Germ Protein Research Kit (WEPRO7240) from CellFree Sciences. The conditions for the synthesis and the 3XFLAG-based purification scheme has been described previously [19].

### 4.3. ProteoLiposome cell-free protein expression of InaZ and TM-InaZ

ProteoLiposome BD Expression Kit from CellFree Sciences has been used. Shortly, 13 μl of pEU InaZ plasmid (1 μg/μl) were mixed with 75 μl of water, 26 μl of 5xTranscription Buffer, 13 μl of 25 mM NTP mix, 1.62 μl of RNase inhibitor and 1.62 μl of SP6 polymerase and incubated at 37°Cfor 5 hours with shaking at 640 rpm on the Thermomixer C incubator from Eppendorf. The entire transcription mixture was further combined with 149 μl of 1xSUB-AMIX SGC buffer, 125 μl of WEPRO7240, 1 μl of 20 mg/ml creatine kinase and 100 μl of freshly hydrated 50 mg/ml Asolectin liposomes (provided as part of Proteoliposome kit). The solution contents were gently mixed by pipetting and transferred to the bottom of the dialysis cup, prefilled with 2 ml of 1xSUB-AMIX SGC buffer. Slide-A-Lyzer Dialysis Device (10K MWCO, 2 ml, ThermoFisher) has been used for the latter where 43 ml of 1xSUB-AMIX SGC were used as a feeding buffer. The translation reaction was carried out in this bilayer-dialysis format overnight at room temperature and in the vibration-free setting. The next day, 2.5 ml of the reaction mixture was gently mixed by pipetting, transferred to a new tube and then centrifuged for 10 min at 13,500g at 4°C. The supernatant was removed, and the pellet was washed three times with 1 ml of TBS buffer using the same centrifugation settings as above. After the final wash, 100 μl of TBS was added to the pellet in order to re-suspend the liposome-protein complexes. The final mixture was split in 10 μl aliquots, which were further flash-frozen in liquid nitrogen and stored at −80°C until future use.

### 4.4. SDS-PAGE electrophoresis

Protein expressions were evaluated on 10% Mini-PROTEAN TGX Precast Protein (Bio-Rad). For sample preparation, 3 μl of a protein solution was mixed with 4 μl of 500 mM DTT and 7 μl of the 2x Laemmi Sample Buffer (Bio-Rad) and then denatured at 90°C for 5 minutes, snap-cooled on ice and loaded on the gel. The general running conditions for SDS-PAGE were 1 hour at 150V in the 1X Tris-Glycine-SDS Running Buffer (Bio-Rad). The gels were further stained with Bio-Safe Coomassie Stain from Bio-Rad. For molecular size estimation and protein yield quantification purposes, Broad Range Protein Molecular Weight Markers from Promega were used.

### 4.5. TEM imaging

An ultrathin carbon film on lacey carbon (01824, Ted Pella) was glow discharged with the carbon side up at 10 mA for 30 seconds using PELCO easiGlow from Ted Pella. Then 3 μl of protein solution (~0.005 mg/ml for InaZ and InaZΔN; 30x dilution for InaZ-liposome complexes) was deposited on the carbon side and let to absorb for 1 minute. The excess of solution was blotted away, and 15 μl of 2% PTA (or NanoW from Nanoprobes) stain has been added on the top and incubation continued for additional 1 min. The excess of the stain was wicked away, and the grid was let to air-dry. Micrographs were acquired on 300kV FEI Environmental TEM.

### 4.6. Ice nucleation experiments using dynamic ESEM

Four samples (1 ml each) were freshly prepared right before the ice nucleation experiments: (1) 0.1xTBS buffer (5 mM Tris-HCl, pH 7.5, 15 mM NaCl, (2) 0.7 μg/ml of InaZΔN in 0.1xTBS buffer, (3) 0.1xTBS buffer supplemented with liposomes, and (4) 12 μg/ml of InaZ-liposome in 0.1xTBS buffer. Samples 1 and 3 were included as negative controls. The tests were also conducted in the low salt buffer to minimize deliquescence. Droplets were generated by nebulization of solution into a nitrogen gas stream at 3 lpm using a medical nebulizer (8900-7-50, Salter Laboratories) and deposited on silicon nitride substrate (Silson Ltd). Experiments were performed in an Environmental Scanning Electron Microscope (Quanta 3D model, FEI, Inc.), equipped with a custom-built ice nucleation platform (IN-ESEM) [31]. The IN-ESEM platform consists of a cryo-cooling stage coupled with a resistively heating element (Minco Product Inc.) and a temperature control unit (Model 22C, Cryogenic Control Systems, Inc.). To monitor the temperature of the droplets, a temperature sensor (±0.15 K, Pt-100, Omega Engineering Inc.) was mounted below the sample holder. A mixing chamber was used to combine ultra-high purity N_2_ (g) from a temperature-controlled water reservoir and dry N_2_ (g) to get desired dew point (T_d_). Humidified N_2_ (g) was then pumped into the ESEM chamber. A chilled mirror hygrometer (GE Sensing, Model 1311XR) was used to measure the T_d_. Imaging was performed using a gaseous secondary electron detector.

Experiments were performed under isothermal conditions where the temperature of the substrate (droplet/particle, T_p_) was kept constant at 250 K, and the partial pressure of water vapor was increased to raise the relative humidity with respect to ice - RH_ice_. Droplets from InaZ-liposome complex were also investigated at multiple temperatures (250K, 240K, 245K, and 235K) at the same final concentration of 12 μg/ml for InaZ. The onset (first nucleation event) was monitored by slow increase in the RH_ice_. A calibration experiment was followed after each ice nucleation event to calibrate T_p_ against T_d_ as described previously [32,33]. The RH_ice_ was calculated based on T_p_ and T_d_ measurements [32,34]. For every sample, three replicate experiments were carried out.

## Acknowledgements

Work was supported by DOE-BER Mesoscale to Molecules Bioimaging Project FWP# 66382. A portion of the research was performed using the Environmental Molecular Sciences Laboratory (EMSL), a national scientific user facility sponsored by the Department of Energy’s Office of Biological and Environmental Research and located at PNNL. We thank Dr. Greg Gloor for assistance in the InaZ sequence correction.

## Author contributions

JEE managed all aspects of this work. JEE and IVN devised experiments for the study. IVN performed genetics, molecular biology, cell-free expression and electron microscopy experiments. SC performed ESEM experiments and data analysis. IVN and JEE wrote initial manuscript draft but all authors contributed to writing the manuscript and approval of final version.

## Supplementary data

DNA and amino acid sequence information for InaZ.

Supplementary Figure Legends.

Fig. S1.

Fig. S2.

Fig. S3.

